# A tool for browsing the Cancer Dependency Map reveals functional connections between genes and helps predict the efficacy and selectivity of candidate cancer drugs

**DOI:** 10.1101/2019.12.13.874776

**Authors:** Kenichi Shimada, Jeremy L Muhlich, Timothy J Mitchison

**Affiliations:** Laboratory of Systems Pharmacology and Department of Systems Biology, Harvard Medical School, Boston, MA 02115, USA

**Keywords:** Cancer Dependency Map, precision medicine, essential genes, CRISPR, RNAi, synthetic lethality, ECHODOTS, cluster analysis, web tool, shiny

## Abstract

Individual cancers rely on distinct essential genes for their survival. The Cancer Dependency Map (DepMap) is an ongoing project to uncover gene dependency in hundreds of cancer cell lines. DepMap is a powerful drug discovery tool, but can be challenging to use without professional bioinformatics assistance. We combined CRISPR and shRNA screening data from DepMap and built a non-programmer-friendly browser (https://labsyspharm.shinyapps.io/depmap) that reports, for each gene, the growth reduction that can be expected on loss of a gene or inhibition of its action (efficacy) and the selectivity of this effect across cell lines. Cluster analysis revealed proteins that work together in pathways or complexes. This tool can be used to 1) predict the efficacy and selectivity of candidate drugs; 2) identify targets for highly selective drugs; 3) identify maximally sensitive cell lines for testing a drug; 4) target hop, *i.e.*, navigate from an undruggable protein with the desired selectively profile, such as an activated oncogene, to more druggable targets with a similar profile; and 5) identify novel pathways needed for cancer cell growth and survival.

## Introduction

Cancer is a disease of the genome. Hundreds, if not thousands, of driver mutations drive cancer in different patients (Bailey et al., 2018), and large collaborative efforts such as the Cancer Genome Atlas Program (TCGA) have helped discover them (The Cancer Genome Atlas Research Network, 2019). Targeted therapies, also called precision medicine, aim to cure cancer by selectively killing cancer cells with a specific genotype and spectrum of driver mutations (Friedman et al., 2015). The underlying hypothesis is that cancers depend on essential genes that are not the same for all cancers, and that these conditionally essential genes constitute a druggable dependency - an “Achilles’ heel” - that can be exploited to develop targeted drugs with minimal toxicity to normal tissues. To achieve this goal, it is important to identify conditionally essential genes for all cancers. It is also important to group conditionally essential genes into related sets, to maximize the chance of finding a druggable target within the set, such as a kinase or other enzyme.

The concept of essential vs non-essential genes arose largely from genetic research in model organisms. Traditionally, it was considered a binary distinction that held across any genotype. However, loss of a given gene can decrease cell growth without killing the cell, so it is more realistic to assign a numerical value to the degree of essentiality, *i.e.*, the extent to which loss of a gene, or inhibition of its product, influences fitness. In cancer, this value may depend on the genotype, transcriptome, and lineage of the cell. In principle, genes that are only essential in a few cell types might make better drug targets, since inhibiting their function is less likely to cause toxicity in non-cancer tissues. For example, the epidermal growth factor receptor is strongly required in certain cancer cells, but not in normal bone marrow stem cells, making it potentially a good target (Wang et al., 2006).

The Cancer Dependency Map project (DepMap) is an ongoing project to identify essential genes across hundreds of cancer cell lines using genome-wide CRISPR and shRNA screens (Tsherniak et al., 2017; Behan et al., 2019). It has already been used successfully to discover genetic vulnerabilities of cancer cells (Sandoval et al., 2018; Wang et al., 2019b). These data represent a gold mine of useful information for biologists and drug developers, but can be challenging for non-bioinformaticians to manipulate and interpret.

The DepMap portal website (https://depmap.org/portal) provides a range of information for each gene, cell lines and lineage dependent on the gene, and co-dependent genes (*i.e.*, other genes whose dependency scores are highly correlated with the gene) as well as basal transcriptome, copy numbers, and mutations of the gene. However, there are no native tools available in DepMap for systematically comparing one gene with all other genes to determine the range of options available for attacking a particular tumor type. We set out to develop a web tool to enable researchers to rapidly determine the essentiality and selectivity of a given gene across cell lines, and to find other genes with similar essentiality profiles.

To explore gene essentiality in the DepMap data we first compared two datasets based on CRISPR and shRNA screens. We found that the degree of concordance was moderate, and that it was useful to combine the data into a unified “perturbation score”. Other recent studies also showed that similar approaches are useful (Gilvary et al., 2019; Wang et al., 2019a). For each gene, this score reports the degree to which loss of the gene reduces cell growth in sensitive lines (“efficacy”), and the degree to which its essentiality varies across lines (“selectivity”). We then visualized all genes on a 2D plot, and defined an essential subset using an arbitrary threshold on efficacy. Next, we clustered perturbation scores of essential genes to discover shared essentiality. Many clusters represent complexes or biological pathways, as previously reported (Pan et al., 2018). We made the analysis accessible as a simple interactive web-tool (https://labsyspharm.shinyapps.io/depmap) to help others identify target genes for anti-cancer drugs, and to discover the mechanistic basis of essentiality in specific cell types.

## Results

### shinyDepMap: an interactive web tool to explore essentiality of genes

shinyDepMap was made utilizing the shiny package of the R language (Chang et al., 2019). It consists of two services, “Essentiality-Selectivity (all genes)” and “Functional clusters (essential genes only)”. shinyDepMap is a dashboard-style website that is split into three components when opened in a web browser: menu bar, input sidebar, and output (Fig. 1A). The services can be opened from the menu bar. This tool can be accessed in two ways, by querying our web site or by downloading the code and pre-processed data from the code repository (https://github.com/kenichi-shimada/shinyDepMap) and run it on a local computer. The first two sources will be maintained and updated by the authors when the original data in the DepMap project are updated. The last method comes with the executable R with all the packages installed, and therefore it is easiest to run the tool locally, but it will not be updated. In the following, the analysis workflow is explained (Fig. 1B).

**Figure 1.**
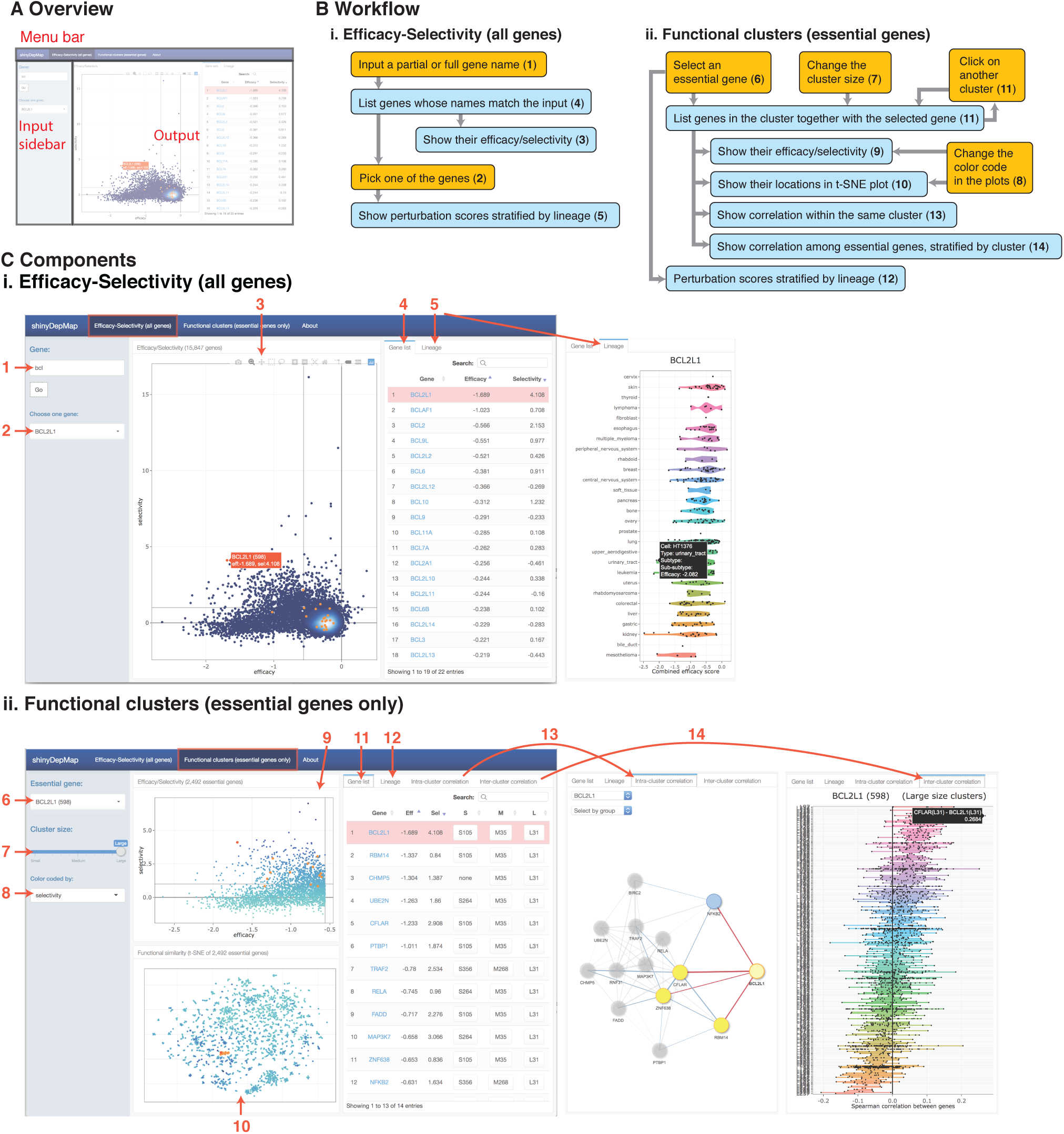
shinyDepMap, an interactive web-tool to explore the DepMap genetic perturbation data. **A. Layout.** There are three parts, menu bar (select analysis), input sidebar (select genes and specify parameters), and output. shinyDepMap lets a user explore the DepMap data in two ways: i) efficacy and selectivity of genes, ii) functional clusters of essential genes. **B. Workflow.** Orange and blue indicate inputs from user and outputs by the website, respectively. Numbers correspond in C. **C. Components in the tool.** **i. Efficacy-selectivity.** 1. Gene symbol query 1. **2.** Gene symbol query 2. **3.** Efficacy-selectivity scatter plot. **4.** Hit gene list. **5.** The perturbational score profiles of a selected gene stratified by lineage. **ii. Functional clusters. 6.** Gene symbol query. **7.** Cluster size (three levels). **8.** Color codes of plots. **9.** Efficacy-selectivity of essential genes. **10.** example t-SNE plot of the perturbation score profiles. **11.** Gene lists of the cluster. **12.** The perturbations score profiles of a selected gene (the same information as 5). **13-14.** Spearman correlation coefficients between the genes within the same cluster (**13**) or across all the clusters (**14**).

#### Efficacy-Selectivity (All genes)

This service allows a user to explore efficacy and selectivity. The output is split into two panels. On the left, a scatterplot displaying efficacy and selectivity of all the genes is always shown (**3**, the number corresponds in Fig. 1B-C). By hovering over the points on the plot with the cursor, one can identify specific genes. When a partial or full gene name is provided in a text box in the input sidebar (**1**), genes that are partially matched to the query as well as their efficacy and selectivity are listed on the right panel, when the “Gene list” tab is selected (**4**). One of the listed genes is highlighted in pink. The points on the scatter plot corresponding to the listed genes are shown in orange with the highlighted gene on the list in red. When switching to the “Lineage” tab, perturbation scores of the highlighted gene in individual cell lines are shown, stratified by lineages (**5**). To view the scores of other genes on the Gene list, the user selects a different gene from the drop-down menu in the Input sidebar (**2**).

#### Functional clusters (essential genes only)

2,492 genes were found essential in our analysis. We clustered perturbation profiles of the genes, as described in detail below. The output consists of three panels. On top left is the same essentiality vs selectivity plot of the essential genes (**9**). The bottom left plot shows the similarity of perturbation score profiles across genes (**10**). Adjacent points in the plot are expected to be functionally connected. When one essential gene is chosen from the top left drop-down menu (**6**), the cluster containing the query gene is listed in the “Gene list” tab on the right output panel (**11**), and the same genes are highlighted in orange and red in the left two plots, **9** and **10**. The clustering was run in three ways. The analyses are different in the size and the tightness of the resulting clusters. The hierarchy is constructed such that tighter/smaller clusters are merged to generate larger/looser clusters. In shinyDepMap, the tightest/smallest clusters are shown by default, but the user can view larger clusters by changing the “Cluster size” slider input (**7**). Alternatively, genes in a larger cluster can be shown by clicking the cluster (S∼/M∼/L∼ for small/medium/large size clusters). The “Lineage” tab shows the perturbation score profiles of the selected gene (**12**), which is the same as (**5**). “Intra-cluster correlation” tab shows the genes in the same cluster as a network representation where two genes are connected when they are positively correlated (Spearman correlation coefficients > 0.1) (**13**). “Inter-cluster correlation” tab shows the Spearman correlation coefficients between the selected gene and all the essential genes, stratified by clusters (**14**). Hovering over a point with the cursor in any scatter plots will give more information, such as a gene name and clusters.

### Computation of a unified ‘perturbation score’

The DepMap website (https://depmap.org/) provides two separate pre-processed genome-wide genetic perturbation data in hundreds of cell lines, in which shRNA and CRISPR efficacy data were normalized using DEMETER2 and CERES algorithms, respectively (Meyers et al., 2017; McFarland et al., 2018). In both, genes with more negative values are considered essential in the corresponding cells. Since the algorithms take ‘off-targets’ into consideration, the normalized CRISPR and shRNA data should give similar scores. To assess this consistency, we compared CRIPSR and shRNA scores in the 4,846,055 equivalent conditions, where the same genes were targeted in the same cell lines (Fig. 2A). We observed a generally high degree of consistency between the two technologies, with some biases. CRISPR tends to detect weak to moderate effects of gene deletion more sensitively while shRNA tends to detect moderate to strong effects (Fig. S1A,B). While CRISPR is claimed to be much less susceptible to off-target effects (Smith et al., 2017), it was most informative to combine CRISPR and shRNA data together considering their different dynamic ranges. We developed a new gene dependency metric, termed “perturbation score”, which is equivalent to a primary principal component of the two efficacy measures (Fig. 2B). The resulting data consist of the perturbation scores of 15,847 genes in 423 cell lines, in which both screenings were performed (Supplemental Data 1).

**Figure 2.**
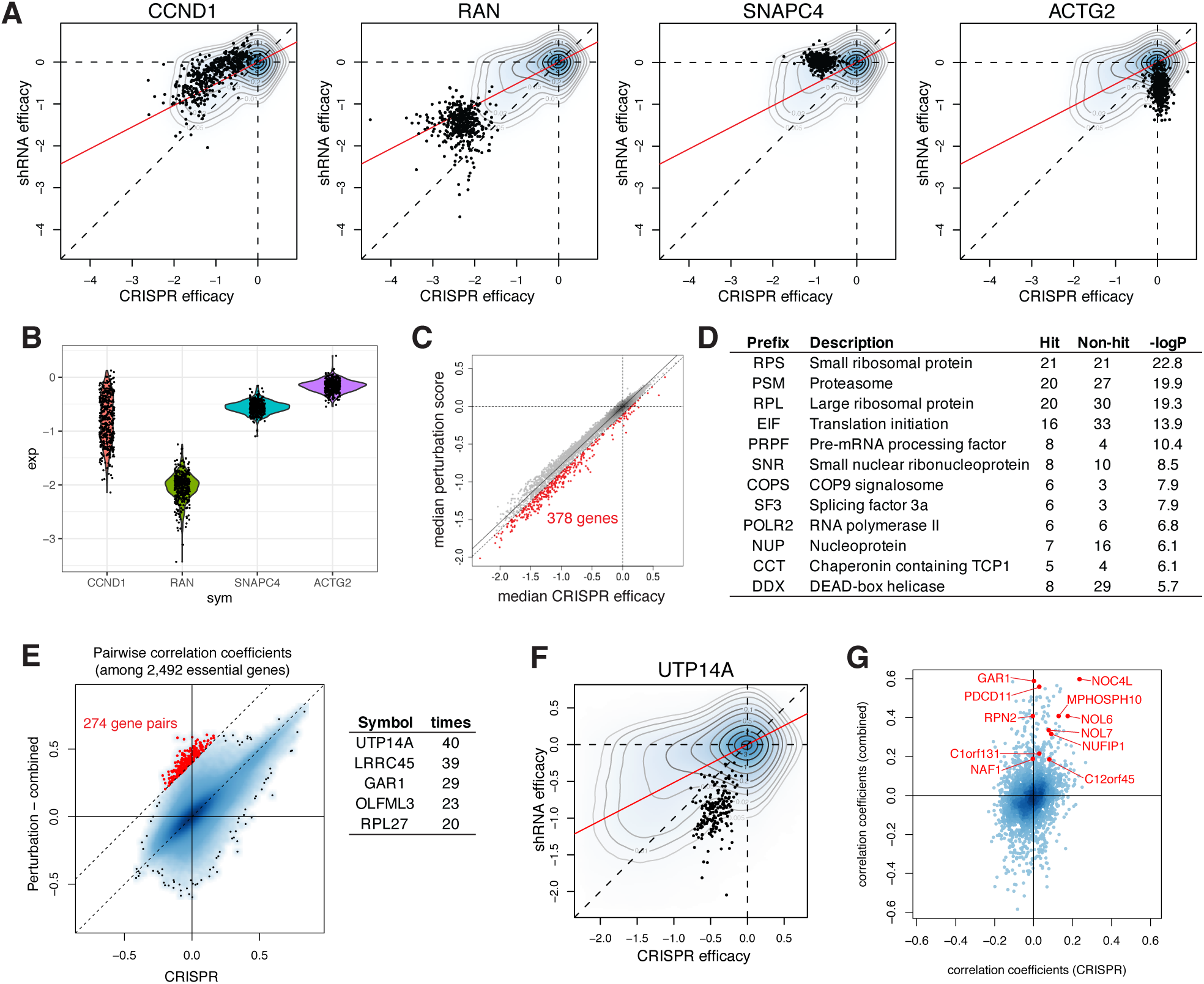
Identification of essential genes across 423 cancer cell lines. **A.** Distribution of CRIPSR and shRNA efficacy scores of 4,846,055 equivalent conditions. Data points indicate CRIPSR/shRNA targeting a gene on the title in all the cell lines. Examples are two consistent (CCND1 and RAN) and two inconsistent (SNAPC4 and ACTG2) genes. Red line indicates the direction of the eigenvector corresponding to the primary principal component. **B.** The perturbation score profiles of the same four genes as in **A**. **C.** Comparison of median CRISPR efficacy and median perturbation score of 15,847 genes. 378 genes in red are the genes whose perturbation scores are lower than CRISPR efficacy by 0.1 or more. **D.** Prefix of gene symbols that are overrepresented in the 378 genes in C. -logP is negative log transformation of p-value computed using Fisher’s exact test. **E.** Spearman correlation coefficients between 2,492 essential genes. Correlation was calculated based on the CRISPR efficacy and perturbation scores, respectively. Correlation coefficients of the 274 gene pairs in red were greater by 0.4 or more when calculated based on perturbation scores than calculated based on CRISPR efficacy. Top five genes frequently seen are listed. **F.** CRISPR and shRNA efficacy targeting UTP14A. **G.** Spearman correlation coefficients between UTP14A and the other essential genes. Genes clustered with UTP14A in the later analysis were highlighted in red. See also **Fig. S1**.

We next compared the median perturbation scores across cell lines for each gene with the median CRISPR and shRNA efficacy. The perturbation scores were much more correlated with CRISPR efficacy than with shRNA efficacy (Fig. 2C, Fig. S1C,D). While the perturbation score and CRISPR efficacy were highly correlated, genes involved in certain biological functions, such as transcription, pre-mRNA processing, translation, chaperonin, and protein degradation, were found to be more essential overall according to the perturbation score over CRISPR efficacy. shRNA-induced knockdown of these genes had larger effects than knockdown of many other genes, for unknown reasons (Fig. 2D). We identified 2,492 essential genes in later analysis and assessed their functional similarity based on the Spearman correlation of their dependency profiles across all the cell lines. The functional similarities between the essential genes computed from the CRISPR or shRNA efficacy alone are not correlated (Fig. S1E). The functional similarities from the perturbation score and the CRISPR efficacy have good agreement, but some gene pairs were found more strongly positively or negatively correlated using the perturbation score while little to no correlations using the CRISPR efficacy (Fig.2E, Fig. S1E). UTP14A, a gene involved in ribosome biogenesis, was one of such genes. The stronger correlations using the perturbation score can be due to the shRNA data since these genes were scored more essential using shRNA than using CRISPR (Fig. 2F). In later analyses, UTP14A was found clustered together with other essential genes involved in ribosomal biogenesis, however, UTP14A and most of these genes were not significantly correlated using CRISPR (Fig. 2G). In this case, the stronger positive correlations only seen using the perturbation score seems to reflect the reality rather than they are artifacts of the analysis. Thus, we concluded that the perturbation score is as good as CRISPR, and for certain genes, represents a better efficacy measure than CRISPR or shRNA alone. We decided to use the perturbation score for the rest of the analysis.

### Identification of essential genes

We next asked which genes were essential. Cancers take various phenotypes and they depend on different essential genes. A gene can be strongly essential in a few cell lines, while most cell lines do not need it for survival. Parametric description of distributions, such as mean or standard deviation, is not an appropriate way to capture such cases. We chose non-parametric quantiles, namely, the perturbation score of 10^th^ most sensitive cell line for each gene as efficacy. The distribution of the efficacy was asymmetric and had a heavy lower tail when the normal distribution was fit, which likely correspond to essential genes in sensitive cell lines. We identified 2,492 essential genes whose efficacy was < −0.560, which is significantly outside of the normal distribution (*p* < 1e-3) (Fig. 3A). The majority of included genes were robustly identified as essential irrespective of definitions of efficacy (*i.e.*, the perturbation scores of 20^th^ or 42^nd^ sensitive cell lines that roughly correspond to 5^th^ and 10^th^ percentiles, instead of 10^th^ sensitive cell line that corresponds to 2.5^th^ percentiles; Fig. S2A-C).

**Figure 3.**
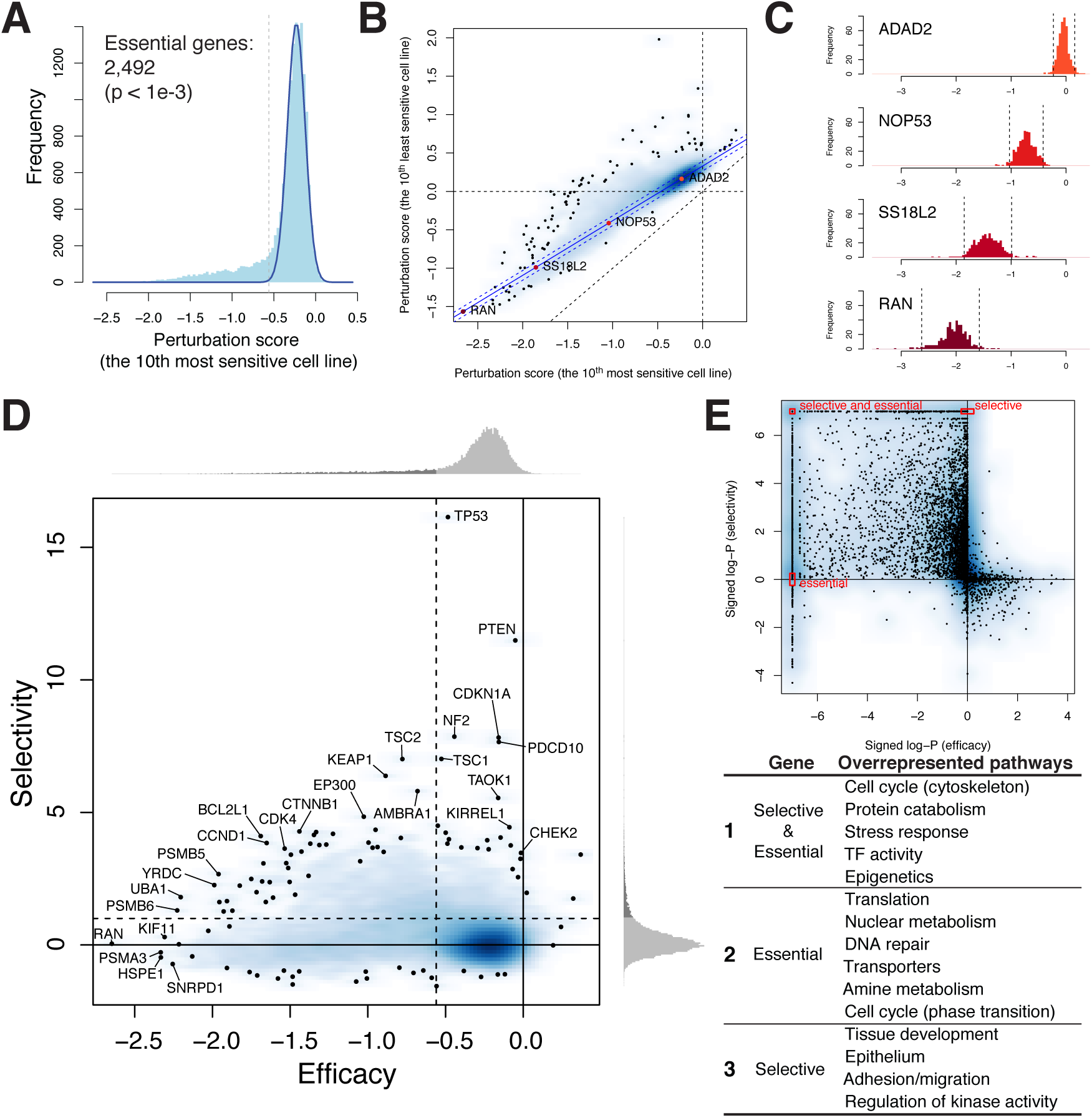
Identification of essential and selective genes. **A.** Distribution of efficacy (perturbation scores of the 10th most sensitive cell lines) among all the genes. Fitted normal distribution is overlaid on top of the distribution. 2,492 genes left to the dashed vertical line were identified as significantly essential by p-value of 1e-3. **B.** Relationship between the perturbation scores of the 10th most and least sensitive cell lines of 15,847 genes. Linear regression was computed using only non-essential genes. Dashed lines parallel to the regression line show the boundary of 95% confidence interval, which contain 80.0% of all the genes. **C.** Distributions of perturbation scores of four genes with different efficacy. The percentile scores of the 10th most and least sensitive cell lines are shown for each distribution. **D.** Efficacy and selectivity of 15,847 genes. **E.** Characterization of efficacy and selectivity in terms of biological pathways. (Top) Pathways significantly overrepresented among essential selective genes give strong negative and positive values on x- and y-axes, respectively. (Bottom) Characteristics of the genes and overrepre-sented pathways in red boxes 1, 2 and 3 of the top panel. See also **Fig. S2, S3**.

### Computation of cell line selectivity

Next, we assessed the selectivity of each gene. The selectivity is an important measure for potential good drug targets for synthetic lethality; drugs targeting generally essential genes may be toxic not only to cancers but can harm other normal cells. Like the efficacy, we non-parametrically defined the selectivity. We first looked at the difference between the perturbation scores of the 10^th^ most and least sensitive cell liens and found that the difference is in a strong linear relationship for about 80% of genes. In other words, the dispersion of the distributions, assessed by the difference is determined by the efficacy (the perturbation score of the 10^th^ most sensitive cells) for these genes. The more essential a gene is, the larger the dispersion is (Fig. 3B, Fig. S2D,E). Taking this relationship into consideration, we defined the selectivity as *s* = (*d* -*d*’)/(*k* * *d*’), where *s* is the selectivity, and *d* and *d’* are the measured and estimated dispersions, and *k=3.772* is a coefficient such that *s* > 1 is considered significantly selective (*p* < 1e-3), when the normality is assumed for the distribution of the selectivity. With this definition, we found 1,309 genes selective, 735 of which were also essential (Supplemental Data 2).

By comparing the efficacy and the selectivity, we found that the genes with high essentiality (*i.e.*, large negative efficacy) tend to be less selective, but selectively essential genes tend to be moderate essential (Fig. 3D). Gene set enrichment analysis of 6,551 existing pathways in Molecular Signature Database (v7.0) revealed that essential and selective genes were characterized by chromatin regulation, for example, essential (but not selective) genes by nuclear metabolism and translation, selective (but not so essential) genes by regulation of kinase activity and tissue development (Fig. 3E, Fig. S3).

### Cluster analysis of the perturbation scores revealed functional modules

The perturbation score across cell lines of a given gene carries information about the cellular context that makes the cell dependent on the activity of that gene. Cluster analysis of these data should reveal pathways that connect genes into functional units. This type of - analysis has several benefits. It may be easier to relate the essentiality of pathways to cancer genotypes than to interpret essentiality for individual genes, and pathway analysis can help identify druggable vulnerabilities at the pathway level that might be missed by single gene analysis. We clustered genes based on the similarity of the perturbation scores across 423 cell lines. For this, we utilized a customized clustering algorithm, termed <u>e</u>nsemble <u>c</u>lustering with <u>h</u>ierarchy <u>o</u>ver <u>D</u>BSCAN <u>o</u>n <u>t-</u>SNE with <u>S</u>pearman distance matrix (ECHODOTS), which extends our previous algorithm (Shimada and Mitchison, 2019) (Fig. 4A). As the name indicates, ECHODOTS relies on t-distributed stochastic neighbor embedding (t-SNE) and subsequently clustering of the coordinates given by t-SNE utilizing density based spatial clustering and noise (DBSCAN) (Maaten and Hinton, 2008; Maaten, 2014; Ester et al., 1996). t-SNE is a non-linear dimensionality reduction algorithm that has become particularly popular in biology to visualize high-dimensional single cell data in 2D. The algorithm compares pairwise distances between all data points both at the original high dimension and at low (two) dimension, and attempts to minimize the difference between the two. t-SNE outperforms many other dimensionality reduction algorithms at displaying data points in 2D in a non-overlapping manner while retaining the local structure of the original data. The resulting low-dimensional plot gathers similar points and isolates dissimilar points, which is a perfect preprocessing of data to be used for certain cluster analyses.

**Figure 4.**
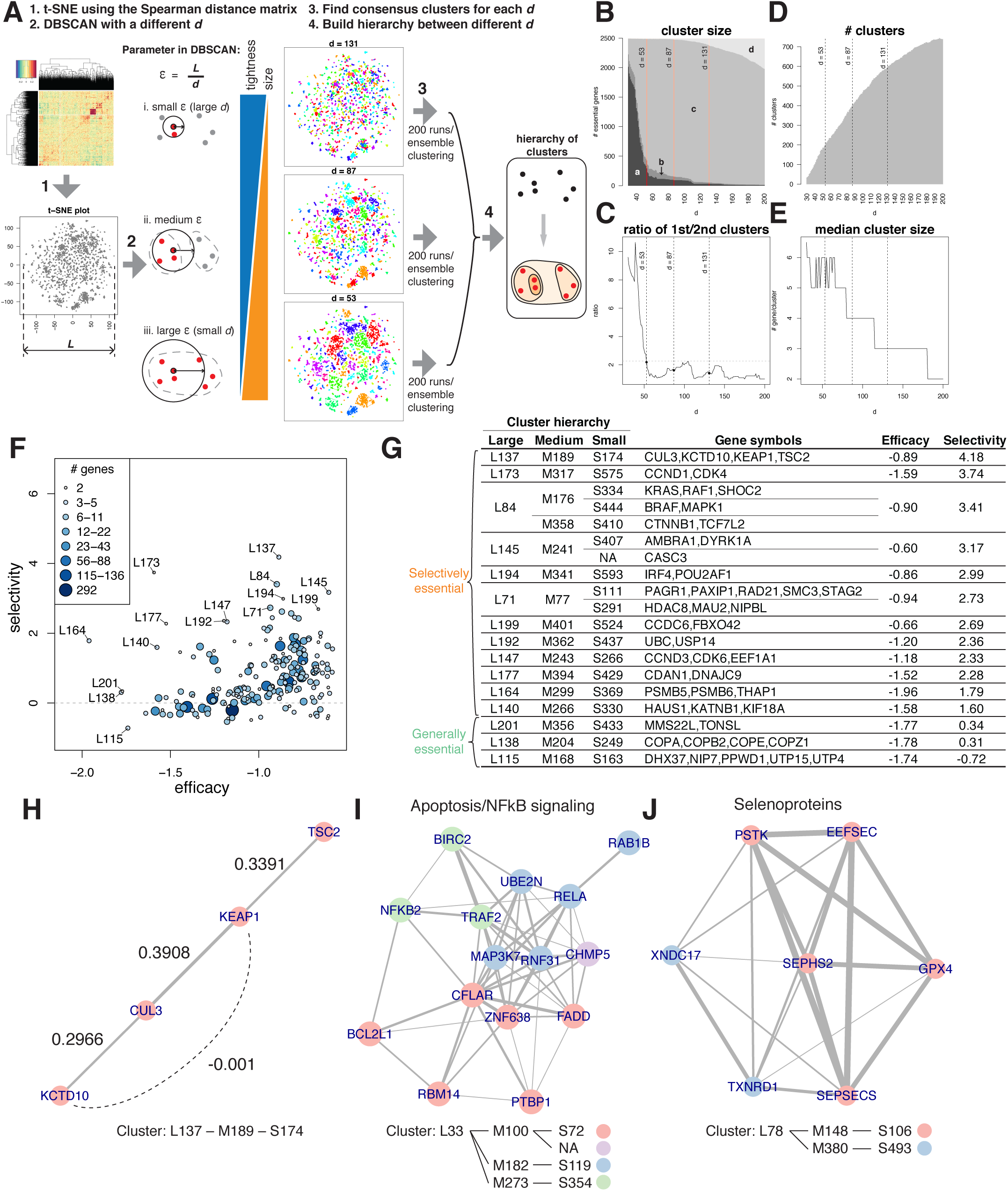
Cluster analysis of essential genes discovered functionally relevant genes together. **A.** Framework of ECHODOTS. See STAR Methods for the detail. **B-E.** Characteristics of identified consensus clusters as a function of d (reciprocally proportional to c). **B.** Cluster sizes. Four colors correspond to genes assigned to the 1st and 2nd largest clusters (**a** and **b**), and other clusters (**c**), or isolated genes (**d**). **C.** Ratio of the sizes of the 1st and 2nd largest clusters. **D.** Number of clusters. **E.** Median cluster size. **F.** Median selectivity and efficacy of the 203 gene clusters (largest cluster sets, corresponding to d = 53). **G.** Clusters that consist of highly selectively essential and generally essential genes. **H-J.** Examples of clusters. **H.** Most selective cluster, L137. **I.** Cluster consisting of anti-apoptosis/NF-kB signal genes. **J.** Cluster consisting of selenoproteins. Genes belong to different tighter/smaller clusters are shown in different colors.

While t-SNE is powerful, it has its own drawbacks. Repeated runs on the same input data produce distinct results due to the non-convexity of the algorithm. This variation between runs may be acceptable for a data visualization purpose as long as they are similar, but robustness can be a serious issue for clustering purposes. To make the cluster analysis more robust, we ran t-SNE and subsequent DBSCAN 200 times and performed ensemble clustering to find genes that are consistently assigned into the same cluster across all runs. In clustering, DBSCAN algorithm takes input data, *i.e.*, t-SNE coordinates, from which distances between any two data points (genes) are calculated, and a neighborhood threshold ε. It assigns two genes into the same cluster when the distance between the genes is smaller than ε. When ε is small, clusters get small and tight. When ε is large, clusters get big and loose. When ε is too large, many data points are connected to become one large cluster (Fig.4B). To avoid over-connection, we set a lower bound for ε (or an upper bound for *d*) such that the size of the 1^st^ and 2^nd^ largest clusters ≤ *b* (*b* = 2.26 in our case) (Fig. 4C). The tighter clusters found with smaller ε are expected to be merged into bigger clusters found with larger ε. We chose three reasonable values of ε, and constructed a hierarchy between the three sets of clusters. We discovered 604 small, 411 medium, and 203 large clusters using ECHODOTS (Fig. 4D, Supplemental Data 3). Some clusters contain as many as 31 genes (small cluster) or 292 genes (large cluster). Many of the clusters has only as few as ≤ 5 genes (83% of the small clusters, 49% of the large clusters). The median cluster size is 6, 4, and 3 genes for small, medium, or large clusters, respectively. Median efficacy and selectivity of the clusters vary widely (Fig. 4F,G).

ECHODOTS clusters essential genes when they are so strongly correlated with each other that they are always found as neighbors to each other on t-SNE map. However, this does not mean that every pair of genes in the same cluster is strongly positively correlated. For a gene to be in a cluster with other genes, the gene is only required to be the neighbor of at least one of them. One should always check the Spearman correlation coefficients to interpret the data. An example of this is cluster L137, which consists of four highly selective essential genes, CUL3, KCTD10, KEAP1, and TSC2 (Fig. 4H). Keap1 and KCTD10 proteins serve as adaptor proteins for Cul3-based E3 ligase. They are each known to bind to the same site of Cul3 to form E3 ligases, but they degrade distinct target proteins (Nrf2 and RhoB, respectively) (Cullinan et al., 2004; Kovačević et al., 2018). Since they bind to the same site on Cul3, they should not bind to Cul3 simultaneously. Indeed, the Spearman correlation coefficients between the four genes revealed that CUL3 is positively correlated with KEAP1 and KCTD10, but KEAP1 is not correlated with KCTD10.

Perhaps more intriguing are clusters that appear to show a connection between specific cellular processes and genes not otherwise known to be involved in that process. One example is cluster L33 (Fig. 4I). This consists of three small clusters. Cluster S72 comprises several genes known to protect cancer cells from apoptosis, such as BCL2L1, CFLAR, and FADD. Clusters S119 and S354 contain genes involved in NF-κB signaling pathway, such as RELA, NFKB2, MAP3K7, and TRAF2. While there is some evidence of a connection between Bcl-xL and NF-κB (Glasgow et al., 2001; Khoshnan et al., 2000), this has not received much attention in cancer research. A network of the correlated genes of this cluster suggests that these genes are particularly well connected through MAP3K7 (encoding TAK1), CFLAR (encoding c-FLIP), and an uncharacterized transcription factor ZNF638. Another example is the cluster L78, which consists of two small clusters (Fig. 4J). Cluster S106 contains an antioxidant enzyme and a selenoprotein GPX4, and the genes involved in selenoprotein synthesis (SEPHS2, SEPSECS, PSTK, EFFSEC) (Squires and Berry, 2008). S493 contains another selenoprotein TXNRD1 and its substrate TXNDC17(Espinosa and Arnér, 2018), which are strongly correlated with four selenoprotein synthesis genes. This cluster likely represents gene that modulate ferroptosis (Abdalkader et al., 2018; Ingold et al., 2018).

## Discussion

Traditional cytotoxic drugs tend to be efficacious, in the sense of killing most cell lines, but not very selective. This translates into treatments that are broad-spectrum but often highly toxic. The goal of precision medicine is to achieve “synthetic lethality” at the cellular level, and thus greatly improve efficacy and selectivity at the organismal level, at a cost of narrower applicability. Major challenges to achieve precision medicine in cancer research include understanding the landscape of essential genes in different cancers and predicting the efficacy and selectivity of new drugs. The DepMap is a powerful repository of data relevant to precision medicine, but difficult for scientists without bioinformatics training to browse. We found that the CRISPR and shRNA genome-wide screening datasets in DepMap were sufficiently concordant that we were able to combine them to generate the ‘perturbation scores’ that were more reliable than data from either one alone. Looking at the 10^th^ most and least sensitive cell lines, we computed ‘efficacy’ and ‘selectivity’ parameters for each gene. Our tool, shinyDepMap, allows the user to rapidly determine the efficacy and selectivity of a gene of interest, and also to find highly selective genes that may offer good targets for precision medicine.

Our cluster analysis of the perturbation scores identified genes that are linked in complexes and pathways as previously reported (Pan et al., 2018). Our analysis complements published work in the area of complex and pathway annotation, in part because we were able to combine both DepMap datasets. We provide cluster information in browsable form in shinyDepMap, including a tool that allows clusters to be made more or less tight. An important use of this tool is “target hopping”, *i.e.*, moving from one drug target to another while keeping the selectivity profile similar (Schenone et al., 2013). One goal in target hopping is to move away from a competitor’s target. Another is to identify druggable targets with similar selectively profiles to genes of interest that are not conventionally druggable. For example, suppose a chemist wanted to target KRAS dependent cancers. KRAS is in cluster L84 with two much more druggable kinase proteins, RAF1 and MAPK1, suggesting that these may be alternative targets based on DepMap data.

In this paper, we only touched on a few findings from our analysis due to limitations of space. shinyDepMap should open up many opportunities for researchers to identify potential druggable cancer targets. However, the interpretation of these data requires much caution. The data in DepMap describes gene requirements for cells growing in monolayer culture in nutrient rich media, a very different environment from the environment of a solid tumor within a patient. Lack of a physiological microenvironment, and in particular the lack of information about the responses of the immune system, may limit clinical translation of gene dependencies. Moreover, the selectivity parameter we developed here is based only on the information on the cell lines included in DepMap, most of which are equivalent to aggressive and malignant cancers and none of which are wild-type non-cancerous. Provided these limitations are kept in mind, DepMap is a powerful resource, and we hope the shinyDepMap tool increases access to it.

## Acknowledgements

Authors thank Laura Maliszewski, Peter Sorger, and Rebecca Ward of Harvard Medical School for helpful discussions. This work is financially supported by Japan Society for the Promotion of Science Overseas Research Fellowship (to K.S.), NIH R35GM131753 (to T.J.M.), and P50GM107618.

## Author Contributions

Conceptualization, K.S. and T.J.M.; Methodology, K.S.; Formal Analysis, K.S.; Software, K.S., T.J.M.; Investigation, K.S.; Data Curation, K.S.; Writing - Original Draft, K.S.; Writing - Review & Editing, T.J.M.; Visualization, K.S.; Supervision, T.J.M.; Funding Acquisition, K.S. and T.J.M.

## STAR Methods

### Data and Code Availability Statement

Following data of the 2019 Q3 release downloaded from the DepMap project website: CRISPR (avana) (“Achilles_gene_effect.csv”), combined RNAi (“D2_combined_gene_dep_scores.csv”), and the cell line metadata (“sample_info.csv”). The CRISPR and shRNA efficacy data were normalized by the Broad Institute. To compute the combined perturbation score, we use the data of 15,847 genes in 423 cell lines, which were examined with both CRISPR and shRNA. The codes generated during this study are available at https://github.com/kenichi-shimada/depmap-analysis (data processing and analysis) and https://github.com/kenichi-shimada/shinyDepMap (standalone shinyDepMap). This study did not generate unique datasets.

### Computation of perturbation scores, a combined effect of shRNA and CRISPR efficacy

4,846,055 conditions that were tested in both CRISPR and shRNA were plotted. Here, one condition is defined as perturbation of one gene either using CRISPR or shRNA in one cell line. We performed principal component analysis on the two datasets. We computed linear combination of the value projected on the line parallel to the eigenvector corresponding to the primary principal components. There were also some missing values in both CRISPR and shRNA datasets. 1,345,642 conditions (20%) were tested using CRIPSR, but not shRNA. 19,001 (0.28%) were tested using shRNA, but not CRISPR. In these cases, the missing values were imputed from the other data using local polynomial regression (Loess) function in R. 4,457 (0.066%) conditions were not tested by either CRISPR or shRNA, which were left as missing. After missing values were imputed, the two values were further combined to compute the perturbation score.

### Similarity between CRISPR and shRNA efficacy and the perturbation scores

To assess the similarity between the three measures (CRISPR, shRNA, or the perturbation score), we first computed each gene’s median dependency across cell lines, which results in representing each gene’s efficacy by one value. We then computed the Spearman correlation coefficients between the three measures. Correlation coefficients between the CRISPR and shRNA, shRNA and the perturbation score, the perturbation score and CRISPR was 0.245, 0.409, and 0.975, respectively. Median CRISPR efficacy and median perturbation score were strongly linearly correlated: *e_p_* = 0.766*e_e_* - 0.006 (R^2^=0.975), where *e_p_*and *e_c_* correspond to the perturbation score and the corresponding CRISPR efficacy. 378 genes were > 0.1 lower than this regression line (*p* < 0.013 using *t*-distribution with 15,845 degree of freedom) (Fig. 2C). meaning their perturbation scores were substantially more negative than their CRISPR efficacy because shRNA efficacy was also more negative. Whether to assess if there is a systematic bias in genes, we looked at the prefix (first three letters) of gene symbols. When the first three letters represent more than one gene family, we added more letters to represent only one class of genes. eg, POL represents RNA polymerases (Pol), but it contains RNA Pol I, II, and III. Only Pol II is overrepresented by the 378 genes here, so POLR2 was chosen instead of POL to make it clear. We computed the enrichment of the gene symbols using one-sided Fisher’s exact test.

The choice of the measure was critical particularly when we assessed the biological functions of the essential genes in our later analysis. We assume genes that work in the same functional unit, *e.g.*, a complex or a pathway, should have similar gene dependency profile, because the unit as a whole is inhibited in the absence of one of the genes. We assessed such functional similarity by computing the same Spearman correlation coefficients between the essential genes. Pairwise Spearman correlation between the 2,492 essential genes was computed for each measure, and the lower triangular matrix of the correlation matrix was compared. While median CRISPR and shRNA efficacy scores were somewhat correlated (0.409), the pairwise correlation coefficients among the essential genes has no correlation between the two (0.079). On the other hand, shRNA and the perturbation score are marginally correlated (0.264), and CRISPR and the perturbation were highly correlated. When looked closely, the perturbation scores tend to give more positive or negative stronger correlations (Fig. 2E). 274 gene-pairs (0.0088%) satisfied the relationship *ρ_p_* > *ρ_e_* + 0.4, where *ρ_p_* and *ρ_c_* corresponds to Spearman correlation coefficients calculated using the perturbation score and CRISPR efficacy, respectively, but we found that a few genes whose shRNA-induced knockdown are effective, highlight the changes.

### ’Efficacy’ parameter and Identification of essential scores

We represented the distribution of perturbation scores of a gene by two parameters, ‘efficacy’ and ‘selectivity’. Efficacy is a measure that indicates how essential a gene is in sensitive cell lines. We computed the perturbation scores of the 10^th^ most sensitive cell lines that represents efficacy of each gene. This roughly corresponds to the 2.5^th^ percentiles among all the cell lines. We next looked at the distribution of efficacy of all 15,847 genes. The distribution has heavy lower tail. We suspected that majority of the gene are non-essential and their deletion has minimal effects on cell growth. Only some genes are expected to be essential in some context. Since there are very few genes whose loss which positively impacts cell growth, we fit the Gaussian curve on the right side of the efficacy’s distribution. The curve was extended to the left-side of the distribution, and genes whose efficacy corresponds to 0.1% of the normal distribution were identified as essential in some context. We also performed the same analysis with different cutoff to call efficacy, namely the perturbation scores of the 20^th^ and 42^nd^ most sensitive cell lines (corresponding to 5^th^ and 10^th^ percentiles), but the majority of the genes were robustly identified as essential.

### ’Selectivity’ parameter for selective gene dependence

We wanted a definitive measure to claim whether a gene is generally or selectively essential. But the relationship between the sensitive scores of the 10^th^ most and least sensitive cell lines revealed that the more essential the gene is, the wider the distribution of their perturbation score was. This linear relationship was also held among non-essential genes. We defined non-essential genes such that they are not identified as essential from the earlier analysis using the perturbation scores of the 10^th^, 20^th^ and 42^nd^ most sensitive cell lines, as well as not identified as promoting its growth using the scores of the 10^th^, 20^th^ and 42^nd^ least sensitive cell lines. We computed linear regression between the scores of the 10^th^ most and least cell lines among non-essential genes and parameters among genes whose loss have minimal effects on cell growth. From the regression, we could compute the expected dispersion between the 10^th^ most and least sensitive cell lines given the efficacy. When the measured dispersion of a gene is higher than this expectation, the gene has a positive selectivity. Thus, we defined the ‘selectivity’ such that *s* = (*d* - *d*’)/(*k* * *d*’), where *s* is the selectivity, and *d* and *d’* are the measured and estimated dispersions. *d’* in the denominator reflects that the distance *d* - *d*’ needs to be scaled by the expected dispersion. *k* is a coefficient to help *s* interpretable: when *s* > 1, it is outside the fitted normal distribution of the selectivity with *p* < 1e-3.

### Pathway analysis to characterize essential and/or selective genes

To characterize biological functions overrepresented among essential genes, selective genes, and essential and selective genes, we performed gene set enrichment analysis (GSEA) utilizing Gene Ontology, KEGG, Reactome, Biocarta, Oncogenic signature, and Hallmark portion of Molecular Signature database (MSigDB) (Subramanian et al., 2005). The MSigDB v7.0 pathway information was downloaded from the Broad Institute (http://software.broadinstitute.org/gsea/msigdb/index.jsp). Statistical significance in GSEA was computed using fgsea package with 10^7^ permutations (Sergushichev, 2016).

### Robust cluster analysis utilizing t-SNE and DBSCAN: ECHODOTS

The goal of our cluster analysis is to find essential genes that form functional units based on their similarity of perturbation profiles across cell lines tested for both CRISPR and shRNA. Our rational is as follows. Some pathways or complexes require all involved genes for the unit to work. Lacking a single gene involved in the unit (*i.e.*, pathway or complex) will demolish its entire function, which cause cell death if the unit is essential. This consequence is the same upon the loss of any genes in the same unit. Thus, genes that form the same functional unit are expected to exhibit similar perturbation score profiles across hundreds of cell lines. The workflow of our cluster analysis is illustrated in Fig. 4A. It consists of four steps: 1. Computing stochastic neighbors of each gene utilizing t-distributed stochastic neighbor embedding (t-SNE) with the Spearman distance matrix. 2. Gene clusters utilizing DBSCAN algorithm with a different neighborhood threshold ε. 3. Run Steps 2-3 for 200 times, and find consensus clusters for each ε. 4. Build hierarchy between consensus clusters across three different ε. In principle, ECHODOTS is a clustering technique relying on t-SNE and DBSCAN. Similar technique was utilized in cluster analysis of high dimensional data, *e.g.*, single-cell RNA-seq data (Mass et al., 2016). We have made a few modifications on this approach to make the most of the stochastic neighbors identified by t-SNE. We termed our method <u>e</u>nsemble <u>c</u>lustering with <u>h</u>ierarchy <u>o</u>ver <u>D</u>BSCAN <u>o</u>n <u>t-</u>SNE with <u>S</u>pearman distance matrix (ECHODOTS).

### Rational using ‘stochastic neighbors’ computed by t-SNE for cluster analysis

At its core, t-SNE computes probability distribution over neighbors for each point from their distance at the original high dimension, and converts it to the probability distribution of the distance in the lower dimension assuming it follows t-distribution with one degree of freedom. The distance in the lower dimension (usually 2- or 3-dimension) is not uniformly transformed from the distance of each pair in high dimension, but it is adjusted based on the probability distribution of the distance for each point such that the number of neighbors are more or less the same for each data point. This number of neighbors is a user-specified parameter, called ‘perplexity’, and we set perplexity = 5. Neighbors determined in t-SNE is therefore called ‘stochastic neighbors’. There is a motivation to utilize stochastic neighbors computed by t-SNE for cluster analysis over neighbors defined in the high dimensional space. In our analysis, we use the Spearman correlation coefficients (ρ_s_) between perturbation score profiles of genes to assess their similarity, or equivalently, we use the Spearman distance (*i.e.*, 1 - ρ_s_) to measure the distances. A ρ_s_ ranges from −1 to 1. One reasonable way to define neighbors is to consider gene pairs whose ρ_s_ is higher than a certain threshold, *e.g.*, 0.3. Only 0.7% of gene pairs among 2,492 essential genes show ρ_s_ > 0.3, which make it reasonable to call the gene pairs neighbors that are functionally connected. However, this definition revealed that the majority of identified genes encode mitochondrial proteins. In fact, MRPL38, a gene encoding protein in a mitochondrial ribosomal subunit, has 186 neighbors. On the other hand, 649 genes have no neighbors. This view does not necessarily comply with our intuition. We often want to know ‘which other genes correlate most with the gene of interest’. For example, CFLAR is the most highly correlated gene with BCL2L1, ρ_s_ is 0.2684. While this is lower than ρ_s_ among most mitochondrial proteins, we are satisfied to call them neighbors since their anti-apoptotic roles are clearly established.

### Cluster ensembles to compute more robust clusters

We next computed clusters utilizing DBSCAN algorithm and feeding the t-SNE output as an input. DBSCAN called a set of genes a cluster when each gene is closer to any other genes than a user-specified threshold ε. We set that each cluster consists of two or more genes. DBSCAN uniquely determines clusters when ε is specified and the input is provided. However, t-SNE does not converge to a unique result as it contains randomness, which is harmful to the cluster analysis. Thus, we ran t-SNE and subsequent DBSCAN 200 times independently with an equivalent ε, and assessed which genes are consistently clustered across the run. Consistent clusters were sought using clue package in R. Note that it was not appropriate to give the constant ε in different runs of DBSCAN since the size of the distribution of t-SNE outputs varied substantially among different runs. Instead, we set ε relative to the size of each distribution (*L*), such that ε = *L*/*d*, where *d* was set constant to give equivalent ε across the runs.

### Different neighborhood threshold gives hierarchy of clusters

Consistent clusters depend on the neighborhood threshold ε. When ε was large (correspondingly, *d* was small), we had a few large clusters. Particularly, the majority of genes formed one huge cluster in an extreme case (d ≤30). When ε got smaller (larger *d*), we got more clusters that were also smaller and tighter. At some point (d ≥ 80), ε became so small that some genes start not having any neighbors. These genes were not in any clusters, since a cluster should have at least one gene. Eventually all the genes becomes isolated so there are no clusters, but we stopped our analysis far ahead (d=200). We chose three different values of *d* (*d* = 53, 87, 131), which discovered roughly 200, 400, and 600 clusters. As expected, smaller *d* (larger ε) gave larger clusters. almost all of which consisted of the smaller clusters from larger *d*. We constructed the hierarchical tree structure between the three sets of clusters. There were only a few genes and clusters that do not fit within this hierarchy. These inconsistent genes that belong to different tree of clusters were flagged as isolated, or manually corrected to support the hierarchy.

### Implementation of shinyDepMap website

shinyDepMap was built using shiny and flexdashboard packages. Following packages are also used to implement the tool: ggplot2 (v3.2.1), RColorBrewer (v1.1-2), shinyWidgets (v0.4.8), plotly (v4.9.0), DT (v0.8), visNetwork (v2.0.8), tibble (v2.1.3), dplyr (v0.8.3), tidyr (v0.8.3). shinyDepMap can be run locally without internet connection. One can download the code and data at https://github.com/kenichi-shimada/shinyDepMap, and run locally following the instruction on the link.

## Supplemental Information

**Figure S1 (Related to Figure 2).**
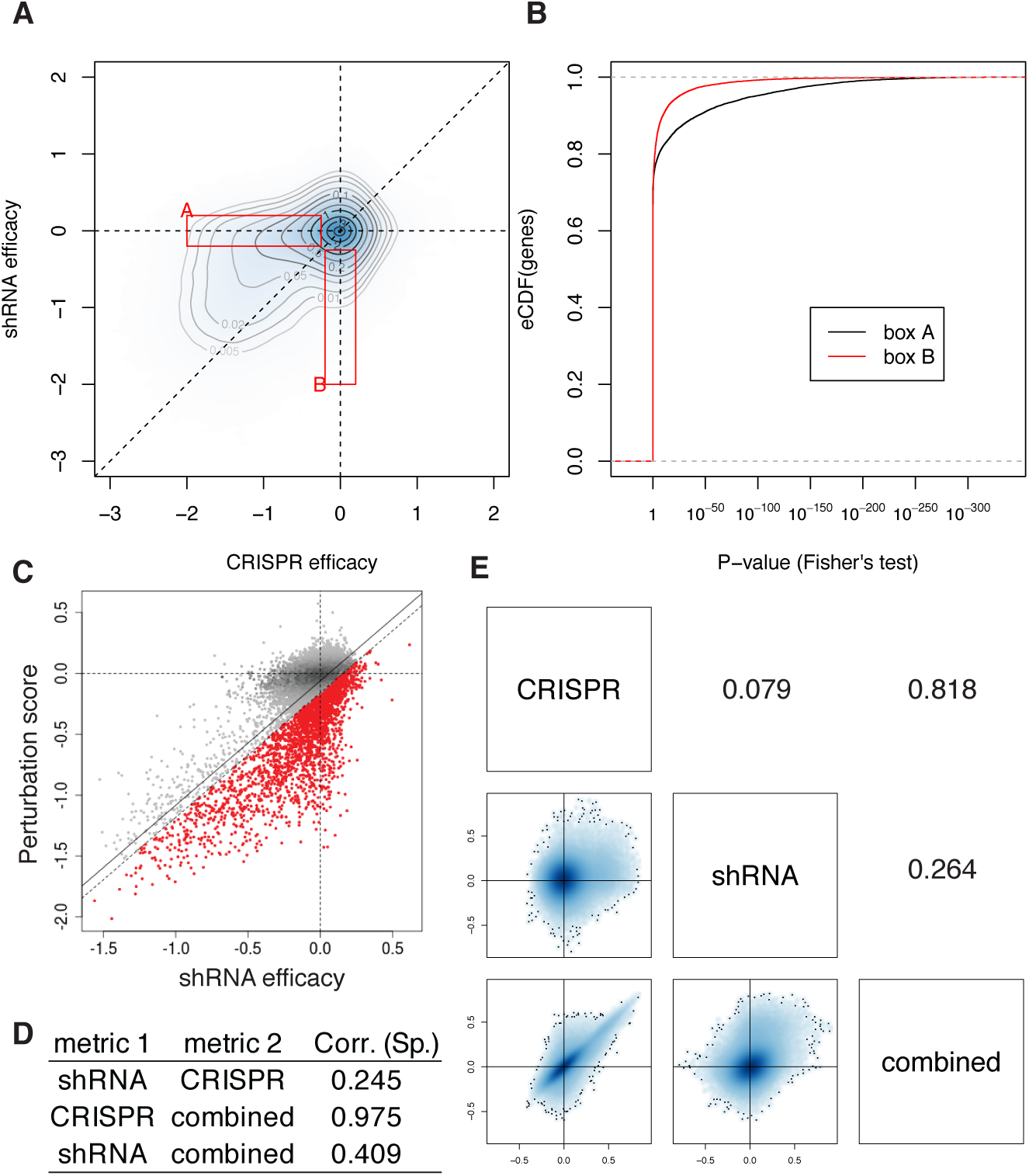
Comparison between CRISPR, shRNA, and the perturbation score. **A.** Search for genes responsive only to CRISPR or shRNA. Red boxes A and B correspond to genes whose suppression by CRISPR or shRNA were found effective to suppress cell growth respectively, but not by the other technology. SNAPC4 and ACTG2 (Fig. 2A) were examples overrepresented in each box. **B.** Overrepresentation of genes in boxes A and B in **A**. Genes overrepresented in each box were assessed by empirical cumulative distrubtion function of p values computed by Fisher’s exact test. **C.** Median efficacy scores using shRNA efficacy and the perturbation score. Median value across the cell lines is plotted for each gene. Red points correspond to the gene whose median shRNA efficacy is lower than the linear regression by 0.1 or more. **D.** The Spearman correlaion coefficients between median efficacy scores for the three measures, shRNA, CRIPSR, and the perturbation score. **E.** Comparison of the Spearman correlation coefficients among 2,492 essential genes, generated using three efficacy measures. Bottom left is the scatter plots, and top right is the corresponding Spearman correlation coefficients between the two measures.

**Figure S2 (Related to Figure 3).**
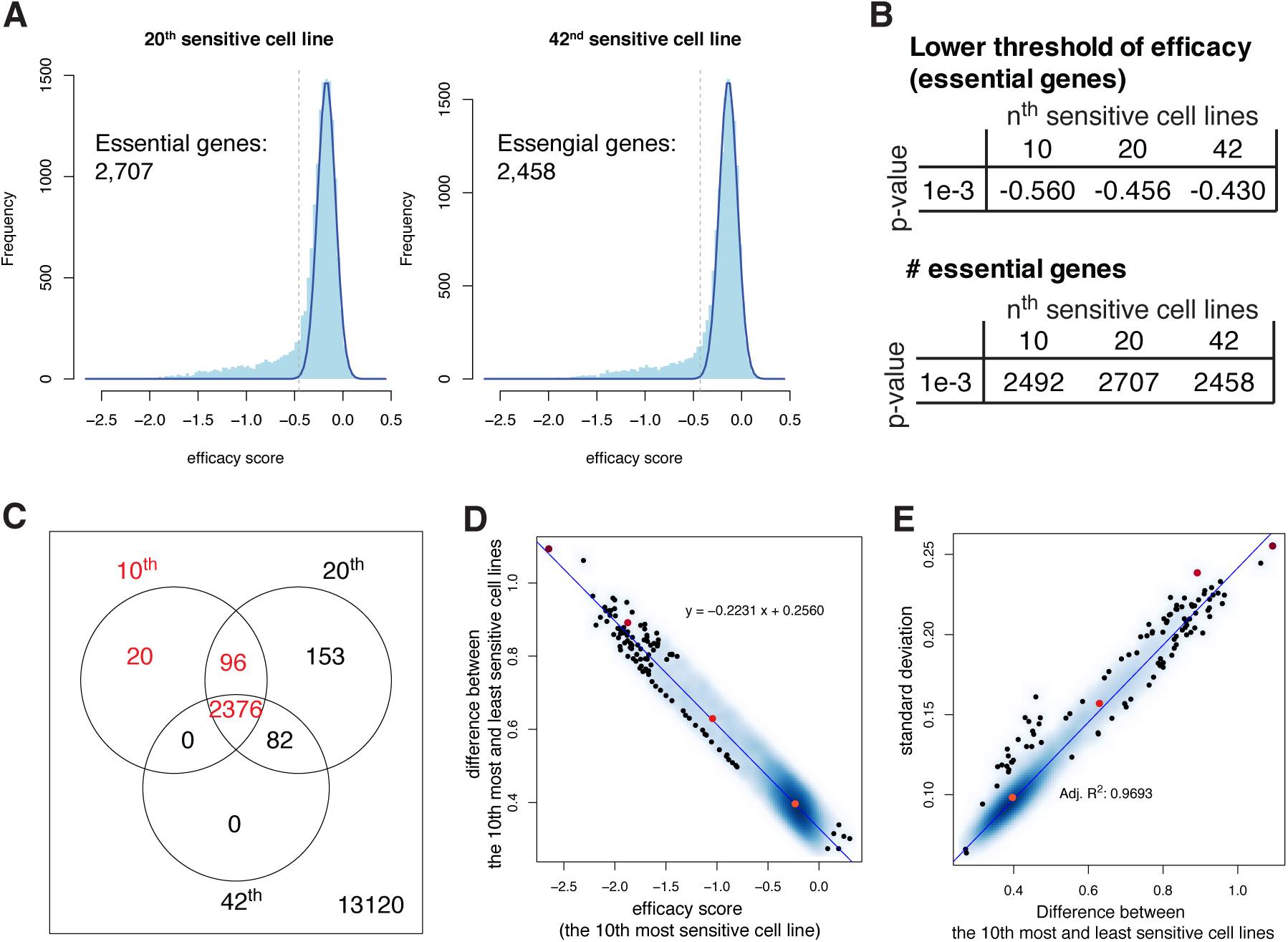
Different cutoffs for defining essential genes. **A.** Distribution of 5th and 10th percentiles of the perturbation scores (corresponding to the perturbation scores of the 20th and 42th most sensitive cell lines). A Gaussian curve was fit on each distribution and to the left of the curves was defined as essential genes. The dashed lines, corresponding to p < 1e-3. **B.** Thresholds (top) and the number of identified essential genes (bottom), using different criteria for efficacy. **C.** Venn diagram indicates the overlap of identified efficacy scores among three definitions of efficacy. **D.** Relationship between the efficacy and the difference between the perturbation scores of the 10th most and least sensitive cell lines. The genes within 95% confidence interval from the regression line in Fig. 3B are shown. **E.** Relationship between the efficacy and the difference between the perturbation scores of the 10th most and least sensitive cell lines.

**Figure S3 (Related to Figure 3).**
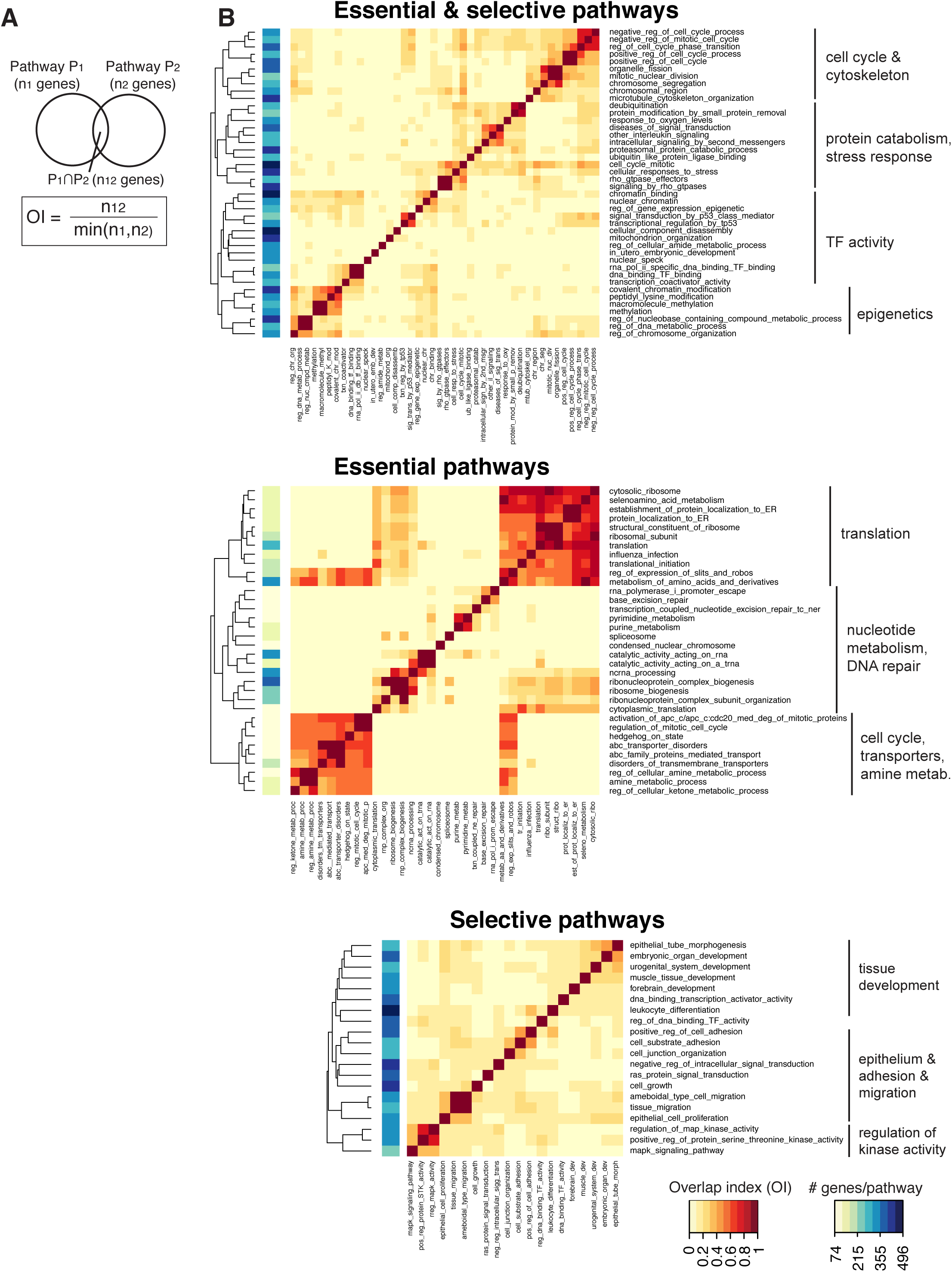
Pathway enrichment against essential and selective genes. **A.** Definition of overlap index (OI) of two pathways, which describes how many genes are overlapped between two pathways. **B.** Overlap index of pathways that are significantly overrepresented in 1) essential and selective genes, essential genes and selective pathways.

**Supplemental Data 1.** The perturbation scores for 15,847 protein coding genes in 423 cell lines

**Supplemental Data 2.** Efficacy and Selectivity for 15,847 proteins

**Supplemental Data 3.** Cluster membership of 2,492 essential genes and the hierarchy of the clusters.

## References

Abdalkader, M., Lampinen, R., Kanninen, K.M., Malm, T.M., and Liddell, J.R. (2018). Targeting Nrf2 to Suppress Ferroptosis and Mitochondrial Dysfunction in Neurodegeneration. Front Neurosci 12.

Bailey, M.H., Tokheim, C., Porta-Pardo, E., Sengupta, S., Bertrand, D., Weerasinghe, A., Colaprico, A., Wendl, M.C., Kim, J., Reardon, B., et al. (2018). Comprehensive Characterization of Cancer Driver Genes and Mutations. Cell 173, 371–385.e18.

Behan, F.M., Iorio, F., Picco, G., Gonçalves, E., Beaver, C.M., Migliardi, G., Santos, R., Rao, Y., Sassi, F., Pinnelli, M., et al. (2019). Prioritization of cancer therapeutic targets using CRISPR-Cas9 screens. Nature 568, 511–516.

Chang, W., Cheng, J., Allaire, J.J., Xie, Y., and McPherson, J. (2019). shiny: Web Application Framework for R.

Cullinan, S.B., Gordan, J.D., Jin, J., Harper, J.W., and Diehl, J.A. (2004). The Keap1-BTB Protein Is an Adaptor That Bridges Nrf2 to a Cul3-Based E3 Ligase: Oxidative Stress Sensing by a Cul3-Keap1 Ligase. Molecular and Cellular Biology 24, 8477–8486.

Espinosa, B., and Arnér, E.S.J. (2018). Thioredoxin-related protein of 14 kDa as a modulator of redox signalling pathways. British Journal of Pharmacology 544–553.

Ester, M., Kriegel, H.-P., Sander, J., and Xu, X. (1996). A Density-based Algorithm for Discovering Clusters a Density-based Algorithm for Discovering Clusters in Large Spatial Databases with Noise. In Proceedings of the Second International Conference on Knowledge Discovery and Data Mining, (Portland, Oregon: AAAI Press), pp. 226–231.

Friedman, A.A., Letai, A., Fisher, D.E., and Flaherty, K.T. (2015). Precision medicine for cancer with next-generation functional diagnostics. Nature Reviews Cancer 15, 747–756.

Gilvary, C., Madhukar, N.S., Gayvert, K., Foronda, M., Perez, A., Leslie, C.S., Dow, L., Pandey, G., and Elemento, O. (2019). A machine learning approach predicts essential genes and pharmacological targets in cancer. BioRxiv 692277.

Glasgow, J.N., Qiu, J., Rassin, D., Grafe, M., Wood, T., and Perez-Pol, J.R. (2001). Transcriptional regulation of the BCL-X gene by NF-kappaB is an element of hypoxic responses in the rat brain. Neurochem. Res. 26, 647–659.

Ingold, I., Berndt, C., Schmitt, S., Doll, S., Poschmann, G., Buday, K., Roveri, A., Peng, X., Porto Freitas, F., Seibt, T., et al. (2018). Selenium Utilization by GPX4 Is Required to Prevent Hydroperoxide-Induced Ferroptosis. Cell 172, 409–422.e21.

Khoshnan, A., Tindell, C., Laux, I., Bae, D., Bennett, B., and Nel, A.E. (2000). The NF-κB Cascade Is Important in Bcl-xL Expression and for the Anti-Apoptotic Effects of the CD28 Receptor in Primary Human CD4+ Lymphocytes. The Journal of Immunology 165, 1743–1754.

Kovačević, I., Sakaue, T., Majoleé, J., Pronk, M.C., Maekawa, M., Geerts, D., Fernandez-Borja, M., Higashiyama, S., and Hordijk, P.L. (2018). The Cullin-3-Rbx1-KCTD10 complex controls endothelial barrier function via K63 ubiquitination of RhoB. J Cell Biol 217, 1015–1032.

Maaten, L. van der (2014). Accelerating t-SNE using Tree-Based Algorithms. Journal of Machine Learning Research 15, 3221–3245.

Maaten, L. van der, and Hinton, G. (2008). Visualizing Data using t-SNE. Journal of Machine Learning Research 9, 2579–2605.

Mass, E., Ballesteros, I., Farlik, M., Halbritter, F., Günther, P., Crozet, L., Jacome-Galarza, C.E., Händler, K., Klughammer, J., Kobayashi, Y., et al. (2016). Specification of tissue-resident macrophages during organogenesis. Science 353, aaf4238.

McFarland, J.M., Ho, Z.V., Kugener, G., Dempster, J.M., Montgomery, P.G., Bryan, J.G., Krill-Burger, J.M., Green, T.M., Vazquez, F., Boehm, J.S., et al. (2018). Improved estimation of cancer dependencies from large-scale RNAi screens using model-based normalization and data integration. Nat Commun 9.

Meyers, R.M., Bryan, J.G., McFarland, J.M., Weir, B.A., Sizemore, A.E., Xu, H., Dharia, N.V., Montgomery, P.G., Cowley, G.S., Pantel, S., et al. (2017). Computational correction of copy number effect improves specificity of CRISPR-Cas9 essentiality screens in cancer cells. Nature Genetics 49, 1779–1784.

Pan, J., Meyers, R.M., Michel, B.C., Mashtalir, N., Sizemore, A.E., Wells, J.N., Cassel, S.H., Vazquez, F., Weir, B.A., Hahn, W.C., et al. (2018). Interrogation of Mammalian Protein Complex Structure, Function, and Membership Using Genome-Scale Fitness Screens. Cell Syst 6, 555–568.e7.

Sandoval, G.J., Pulice, J.L., Pakula, H., Schenone, M., Takeda, D.Y., Pop, M., Boulay, G., Williamson, K.E., McBride, M.J., Pan, J., et al. (2018). Binding of TMPRSS2-ERG to BAF Chromatin Remodeling Complexes Mediates Prostate Oncogenesis. Molecular Cell 71, 554–566.e7.

Schenone, M., Dancfk, V., Wagner, B.K., and Clemons, P.A. (2013). Target identification and mechanism of action in chemical biology and drug discovery. Nat Chem Biol 9, 232–240.

Sergushichev, A. (2016). An algorithm for fast preranked gene set enrichment analysis using cumulative statistic calculation. BioRxiv 060012.

Shimada, K., and Mitchison, T.J. (2019). Unsupervised identification of disease states from high-dimensional physiological and histopathological profiles. Molecular Systems Biology 15, e8636.

Smith, I., Greenside, P.G., Natoli, T., Lahr, D.L., Wadden, D., Tirosh, I., Narayan, R., Root, D.E., Golub, T.R., Subramanian, A., et al. (2017). Evaluation of RNAi and CRISPR technologies by large-scale gene expression profiling in the Connectivity Map. PLOS Biology 15, e2003213.

Squires, J.E., and Berry, M.J. (2008). Eukaryotic selenoprotein synthesis: Mechanistic insight incorporating new factors and new functions for old factors. IUBMB Life 60, 232–235.

Subramanian, A., Tamayo, P., Mootha, V.K., Mukherjee, S., Ebert, B.L., Gillette, M.A., Paulovich, A., Pomeroy, S.L., Golub, T.R., Lander, E.S., et al. (2005). Gene set enrichment analysis: A knowledge-based approach for interpreting genome-wide expression profiles. Proceedings of the National Academy of Sciences of the United States of America 102, 15545–15550.

The Cancer Genome Atlas Research Network (2019). The Cancer Genome Atlas Program.

Tsherniak, A., Vazquez, F., Montgomery, P.G., Weir, B.A., Kryukov, G., Cowley, G.S., Gill, S., Harrington, W.F., Pantel, S., Krill-Burger, J.M., et al. (2017). Defining a Cancer Dependency Map. Cell 170, 564–576.e16.

Wang, W., Malyutina, A., Pessia, A., Saarela, J., Heckman, C.A., and Tang, J. (2019a). Combined gene essentiality scoring improves the prediction of cancer dependency maps. EBioMedicine.

Wang, X., Wang, S., Troisi, E.C., Howard, T.P., Haswell, J.R., Wolf, B.K., Hawk, W.H., Ramos, P., Oberlick, E.M., Tzvetkov, E.P., et al. (2019b). BRD9 defines a SWI/SNF sub-complex and constitutes a specific vulnerability in malignant rhabdoid tumors. Nat Commun 10, 1–11.

Wang, Y., Probin, V., and Zhou, D. (2006). Cancer therapy-induced residual bone marrow injury-Mechanisms of induction and implication for therapy. Curr Cancer Ther Rev 2, 271–279.

